# MxB restricts HIV-1 by targeting the tri-hexamer interface of the viral capsid

**DOI:** 10.1101/444067

**Authors:** Sarah Sierra Smaga, Chaoyi Xu, Brady James Summers, Katherine Marie Digianantonio, Juan Roberto Perilla, Yong Xiong

## Abstract

Myxovirus resistance protein B (MxB) is an interferon-inducible restriction factor of HIV-1 that blocks nuclear import of the viral genome. Evidence suggests that MxB recognizes higher-order interfaces of the HIV capsid lattice, but the mechanistic details of this interaction are not known. Previous studies have mapped the restriction activity of MxB to its N-terminus encompassing a triple arginine motif ^11^RRR^13^. Here we demonstrate a direct and specific interaction between the MxB N-terminus and helical assemblies of HIV-1 capsid protein (CA) using highly purified recombinant proteins. We performed thorough mutagenesis to establish the detailed molecular requirements for the CA interaction with MxB. The results map MxB binding to the interface of three CA hexamers, specifically interactions between positively charged MxB N-terminal residues and negatively charged CA residues. Our crystal structures show that the CA mutations affecting MxB interaction and restriction do not alter the conformation of capsid assembly. In addition, 30 microsecond long all-atom molecular dynamics (MD) simulations of the complex between the MxB N-terminus and the HIV CA tri-hexamer interface show persistent MxB binding and identify a MxB-binding pocket surrounded by three CA hexamers. These results establish the molecular details of the binding of a lattice-sensing host factor onto HIV capsid, and provide insight into how MxB recognizes HIV capsid for the restriction of HIV-1 infection.

**Author summary:** The human antiviral protein MxB is a restriction factor that fights HIV infection. Previous experiments have demonstrated that MxB targets the HIV capsid, a protein shell that protects the viral genome. To make the conical shaped capsid, HIV CA proteins are organized into a lattice composed of hexamer and pentamer building blocks, providing many interfaces for host proteins to recognize. Through extensive biochemical and biophysical studies and molecular dynamics simulations, we show that MxB is targeting the HIV capsid by recognizing the region created at the intersection of three CA hexamers. We are further able to map this interaction to a few CA residues, located in a negatively-charged well at the interface between the three CA hexamers. This work provides detailed residue-level mapping of the targeted capsid interface and how MxB interacts. This information could inspire the development of capsid-targeting therapies for HIV.

## Introduction

The Myxovirus resistance (Mx) proteins are dynamin-like GTPases with antiviral activity. The two Mx proteins found in humans, MxA and MxB, originated from an ancient gene duplication. MxA is a well-characterized restriction factor of influenza and a wide range of other viruses (reviewed in [1]). MxB was recently identified as an interferon-inducible restriction factor of HIV-1 and herpesviruses, as well as murine cytomegaloviruses (MCMV) [2-7]. In cells, MxB localizes to the nuclear periphery, potentially near nuclear pores [8, 9]. MxB appears to restrict HIV approximately 10-fold, and block infection between the steps of reverse transcription and integration [5-7, 9].

MxB is similar to MxA in both sequence and structure but with different viral restriction activities. Each of the Mx proteins is composed of three conserved domains: the stalk domain, the GTPase domain and the bundling signaling element (BSE) domain. The stalk domain is responsible for the oligomerization of Mx proteins. In MxA (and MxB restriction of herpesviruses and MCMV), the GTPase domain is essential for restriction activity [2-4, 6, 7]. The BSE transmits information between domains for viral inhibition [10]. The atomic resolution structures of MxA and MxB [10-12] have shed light on the overall architecture of Mx proteins. The structure of individual monomers is very similar, but substantial differences occur in the higher order dimerization and oligomerization of MxA and MxB [10, 11] Furthermore, MxB contains an N-terminal tail (43 amino acids) that is absent in MxA, and this N-terminus is missing or disordered in the available MxB structures. The unstructured N-terminus of MxB is essential for its restriction activity. Deleting the N-terminal region of MxB, or replacing it with the shorter N-terminus of MxA, abolishes MxB’s restriction activity against HIV-1 [11, 13-15]. Importantly, the restriction activity of MxB can be abolished by mutating a triple arginine motif ^11^RRR^13^ to ^11^AAA^13^ [14].

The HIV-1 restriction activity of MxB is correlated to its ability to bind the viral capsid. It has been reported that certain mutations in the HIV-1 capsid protein (CA) reduce the potency of MxB restriction, suggesting that capsid plays a role in the restriction activity of MxB [5-7, 18]. We and others observed that MxB binds to capsid *in vitro* by using helical assemblies of purified CA and in cells by using the viral capsid core [11, 13, 19]. However, MxB does not bind to the individual capsid building blocks: CA dimers, hexamers, or pentamers [11, 13, 19]. This suggests that MxB is recognizing a feature present only in the higher-order CA lattice on capsid, potentially at an interface between hexamer building blocks. Further evidence to support this hypothesis was provided in an *in vitro* evolution experiment that found two capsid mutations located in the interface between three capsid hexamers, G208R and T210K, that escaped MxB restriction. Other mutations, including P207S, M10I, and P90T, and M185L escaped MxB restriction to lesser degrees [9]. However, the direct role of these residues in CA-MxB interactions is not clear. These experiments were performed by passaging the virus in cell culture, where a capsid mutation that generates a defect in assembly, stability, or interaction with cofactors could affect viral sensitivity to MxB indirectly.

In the present work, we describe how MxB recognizes the HIV capsid by defining the MxB-CA binding interface. Using capsid co-pelleting assays with highly pure MxB and CA components, we performed comprehensive mutagenesis screening of capsid residues that potentially interact with MxB. Our data suggest that MxB recognizes the interface between three capsid hexamers, particularly through interactions between its basic ^11^RRR^13^ motif and acidic CA residues. Our crystal structures show that the MxB-resistant mutations at the tri-hexamer interface do not affect the structures of CA hexamers or the capsid lattice, though they do reduce MxB binding. In addition, results from long time-scale MD simulations demonstrate binding of MxB to the tri-hexamer interface of higher-order capsid assemblies. Furthermore, our simulations establish key residues in CA that mediate interactions with MxB’s N-terminal domain, and are in agreement with the biochemical and biophysical experiments.

## Results

### The N-terminus of MxB binds directly to CA

We produced a minimal MxB construct with good solution behavior and containing the established capsid-binding motif. Previous *in vitro* studies of MxB binding were limited by low yield and poor behavior of full-length MxB protein. Inspired by reports that the N-terminus of MxB dictates restriction and includes the positively charged RRR residues that potentially target capsid [11, 14, 19, 20], we focused our study on the N-terminal 35 residues of MxB. To mimic the extended dimeric architecture of the native protein, which has the two N-termini located at the opposite ends, we fused the N-terminal 35 residues of MxB (MxB_1-35_) to maltose binding protein (MBP). The MxB_1-35_-MBP fusion is further linked to a GCN4 dimerization helix to enforce a dimeric geometry (Fig 1A). The MBP acts as a spacer, creating a separation between the two MxB N-termini, and at the same time serves as a solubility enhancer. This construct, MxB_1-35_-MBPdi, provides a similar geometry to the extended MxB dimer, while allowing us to focus our study on the interactions of its N-terminus without the solution behavior complications of full-length MxB protein.

**Figure 1:**
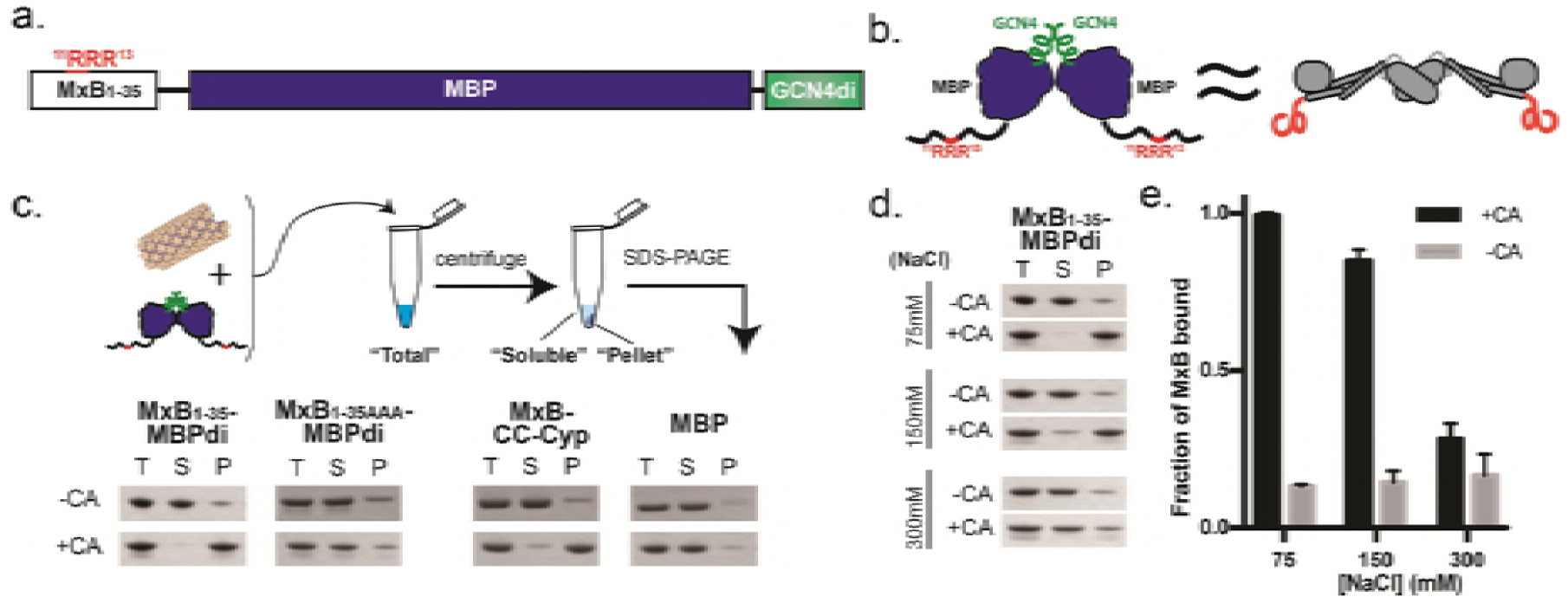
An MxB-MBP dimer construct binds CA helical tubes in vitro. a. Construct schematic of MxB_1-35_-MBPdi.
b. Cartoon model of MxB_1-35_-MBPdi (left), which mimics the extended dimer architecture of the full-length MxB protein (right).
c. Top: Diagram of the co-pelleting assay. Purified CA tubes and MxB constructs are incubated (“total”);after centrifugation, “soluble” and “pellet” fractions are analyzed. Bottom: MxB_1-35_-MBPdi binds CA tubes. MxB_1-35AAA_-MBPdi, in which the RRR CA-binding motif is mutated to AAA binds significantly less. MBP and MBP-CCCyp are used as negative and positive controls, respectively.
d. MxB_1-35_-MBPdi binding to CA tubes is salt-dependent and is abolished at above 300mM NaCl.
e. Quantification of the co-pelleting assay in (d). The fraction of MxB bound was computed for each sample by subtracting the soluble from the total and normalized to the total. Error bars represent the standard error of the mean from three independent experiments.

The MxB_1-35_-MBPdi construct binds directly to CA tubes *in vitro* in a co-pelleting assay, where a soluble capsid cofactor is found in the pellet only upon binding to insoluble polymerized CA [21-23]. We chose to use A14C/E45C disulfide-crosslinked CA tubes in our assay, which mimic the lattice seen in the capsid and tolerate a range of experimental conditions [24]. To quantify MxB binding, we compared the reduction of MxB in the soluble fraction, instead of the amount pelleted, relative to the total input. This approach measures binding at equilibrium and produces more accurate quantification than attempting to measure the MxB bound to the washed or unwashed CA pellet. The results show that over 90% of MxB_1-35_-MBPdi co-pellets with CA at 3 μM MxB concentration, indicating efficient binding under the experimental conditions and validating our MxB construct, which may be useful in future studies of MxB-capsid interactions (Fig 1B). The negative control, MBP alone, has a background level of less than 10% binding. As expected, MxB_1-35AAA_-MBPdi, in which the triple arginine motif known to bind CA is mutated to ^11^AAA^13^, showed much weaker binding than wild-type at 20%. This further validates the construct and the residual binding may indicate that additional residues outside of ^11^RRR^13^ play a minor role in capsid interaction.

The binding of MxB_1-35_-MBPdi to CA tubes decreases upon increasing salt concentration, suggesting that the interaction is driven by electrostatic interactions. At 75mM NaCl, over 90% of MxB_1-35_-MBPdi binds to CA tubes (Fig 1C and 1D). At the near-physiological salt concentration of 150mM NaCl, MxB_1-35_-MBPdi retains around 80% of binding. However, at above 300mM NaCl, the binding of MxB_1-35_-MBPdi is reduced to the ~20% residual level of binding observed in MxB_1-35AAA_-MBPdi. The low-level binding of MxB_1-35AAA_-MBPdi is consistent within error across concentrations of NaCl, suggesting that any remaining interactions may not be charge-driven. To investigate the intrinsic interaction property and more easily distinguish potential subtle mutagenesis effects, we performed subsequent assays at 75mM NaCl, where binding is at its maximum.

### MxB interacts with CA at the interface between three hexamers

We predicted that the positively-charged ^11^RRR^13^ motif of MxB would interact with negatively-charged regions of the CA lattice surface based on its charge character and our observation that MxB binding to CA tubes is salt-dependent. We identified three regions of negative charge on the CA lattice from its calculated surface electrostatic potential distribution (Fig 2A). The residues that contribute to these negatively charged regions include: E71, E75, E212 and E213 at the tri-hexamer interface, E180 and E187 at the di-hexamer interface, and E98 in the CypA-binding loop (Fig 2B). Furthermore, based on the previous observation that individual CA hexamers are not sufficient to confer MxB binding, we posit that the tri-hexamer or di-hexamer interface residues are important for MxB binding. To test these hypotheses, we mutated each of these glutamates, as well as residues in the CypA-binding loop, to alanine and tested their ability to interact with MxB_1-35_-MBPdi in our co-pelleting assay. Our binding results pinpoint the MxB-interacting site at the tri-hexamer interface.

**Figure 2:**
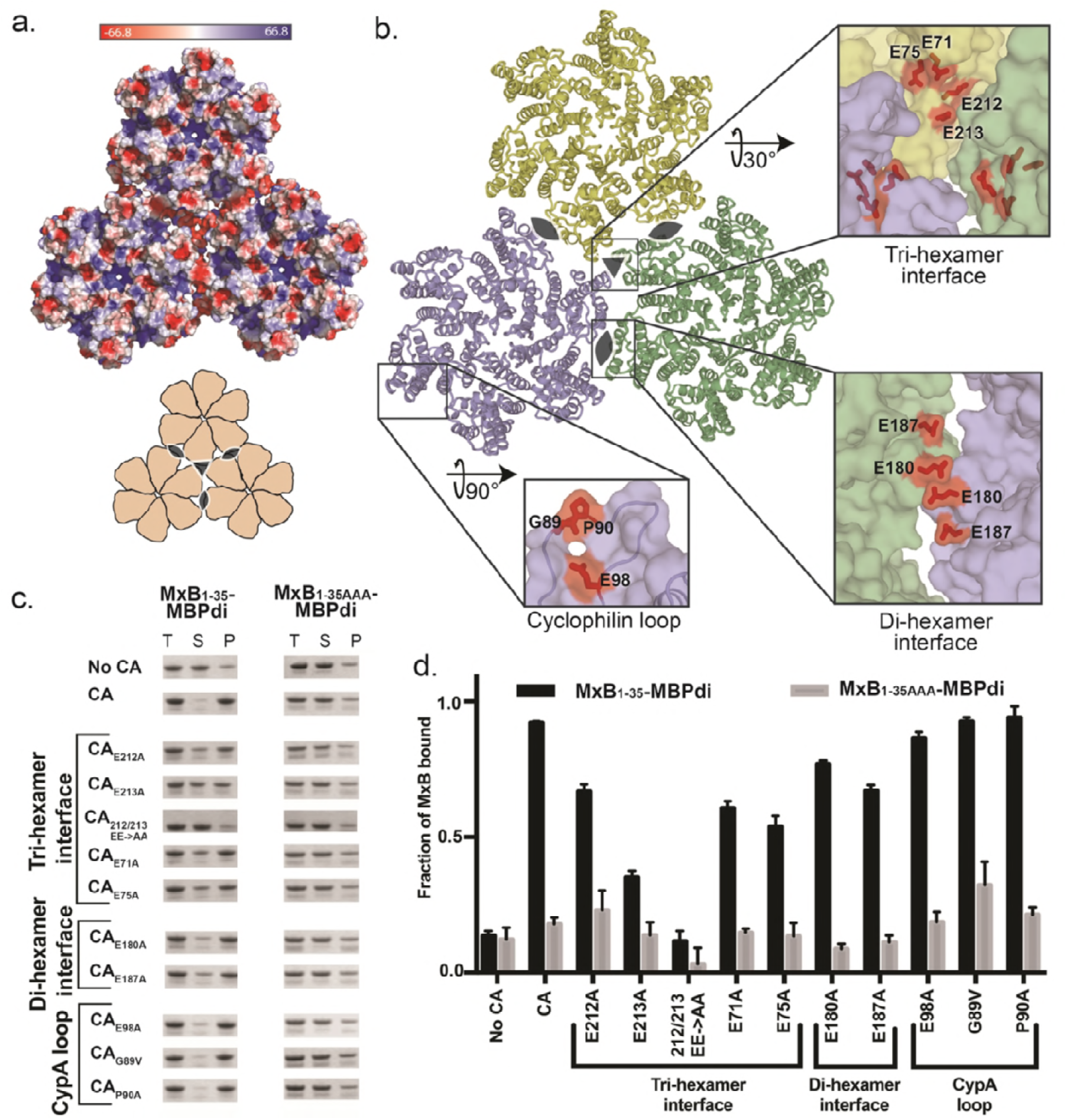
The MxB binding site on capsid maps to the tri-hexamer interface. a. Surface electrostatic potential distribution of three CA hexamers (PDB ID: 4XFX) shows the negative charge at the interfaces between the hexamers (major), and in the CypA-binding loop (minor). A cartoon (bottom) shows the overall arrangement of hexamers and the symmetry axes at the di-hexamer (cat eyes) and tri-hexamer (triangle) interfaces.
b. Location of CA residues selected for mutation. CA hexamers (surface) are shown in light blue, light yellow, and light green. Residues selected for mutation are shown in red sticks.
c. MxB_1-35_-MBPdi binding to CA tube variants in co-pelleting assays. The total, soluble, and pellet fractions were analyzed by SDS-PAGE. For simplicity, only the MxB_1-35_-MBPdi band is shown. MxB_1-35AAA_-MBPdi is used as a negative control. A notable reduction in MxB binding is observed for CA_E213A_, CA_E71A_, CA_E75A_, and CA_212/213 EE->AA_. Positive and negative controls performed with MBP-CCCyp and MBP, respectively, can be found in Supplemental Appendix 1. Full gels can be found in Supplemental Appendix 1.
d. Quantification of data in (c), performed as in Figure 1. Error bars represent the standard error of the mean from three independent experiments. Quantification of controls can be found in Supplemental Appendix 1.

Mutations in tri-hexamer interface residues had the greatest effect on MxB binding. This region of the CA lattice contains the densest concentration of negative charge, with 12 glutamate residues (three copies of E71, E75, E212 and E213) creating a deep well of negative charge. In the tri-hexamer interface, three sets of CA CTD residues E212 and E213 from three adjacent symmetry-related monomers form the floor of this negatively charged well. MxB binding to CA_E213A_ was significantly reduced, and CA_E212A/E213A_ drastically reduced binding to the background level (Fig 2C and 2D). The CA NTD residues E71 and E75 form the wall of the negatively charged well and mutations at these sites (CA_E71A_ and CA_E75A_) also substantially reduced MxB binding (Fig 2C and 2D).

Mutation of residues in the di-hexamer interface has a mild effect on MxB binding. This interface is present in the WT CA dimer, which showed no binding to MxB in solution in previous experiments [11]. We identified glutamates E180 and E187 on helix 9 at the CTD-CTD dimerization interface as responsible for the patch of negative charge between two CA hexamers. Both CA_E180A_ and CA_E187A_ modestly affected MxB binding (Fig 2C and 2D). This suggests they do not contribute substantially to the MxB interaction.

In contrast to the effect from perturbations at the hexamer interfaces, mutation of residues around the CypA-binding loop on the surface of the CA hexamer had no effect on MxB binding. The most prominent region of negative charge on the hexamer surface is contributed by residue E98 near the CypA-binding loop, which is not near any lattice interfaces. Consistent with our interface-binding hypothesis, MxB_1-35_-MBPdi bound CA_E98A_ tubes at a level similar to CA tubes (Fig 2C and 2D). We additionally examined the interactions of MxB_1-35_-MBPdi with CA tubes having mutations in the CypA-binding loop. Since the G89V and P90A mutations have been reported to affect MxB restriction *in vivo* and G89V has been reported to reduce MxB copelleting of MxB in cell lysates with CANC tubes [6, 7, 13], we tested these two mutations in our *in vitro* system using purified MxB and CA components. Our previous work demonstrated that G89V had no effect on full-length MxB pelleting [11]. Using our minimal MxB-MBPdi construct, we confirmed that neither CA_G89V_ nor CA_P90A_ reduce the binding (Fig 2C and 2D).

### Tri-hexamer interface mutations that escape MxB restriction also abolish MxB_1-35_-MBPdi binding

Three mutations at the tri-hexamer interface (P207S, G208R and T210K) (Fig 4A) were previously identified in an *in vitro* evolution experiment to escape MxB restriction to varying degrees [9]. The mutation P207S is relatively conservative, while both G208R and T210K introduce bulky, positively charged residues to the predominantly negatively charged environment described above (Fig 2A, 3A). Based on our interaction mapping results, we predicted that both G208R and T210K, which drastically alter the electrostatic environment of the tri-hexamer interface, would decrease MxB binding to capsid. To test this hypothesis, we introduced these mutations into CA tubes and performed co-pelleting assays with MxB_1-35_-MBPdi.

The CA mutations G208R and T210K drastically reduced MxB binding (Fig 3B). MxB_1-35_-MBPdi binding to CA_G208R_ was indistinguishable from the MxB_1-35-AAA_-MBPdi background signal. Binding to CA_T210K_ was also substantially reduced. By neutralizing the strong negative charge at the tri-hexamer interface, these mutations likely disrupt charge-charge interactions that are required for MxB binding. It is also possible that the bulkiness of these residues occludes the well at the tri-hexamer interface and sterically blocks MxB access. The relatively modest mutation P207S had a minor effect on MxB interaction (Fig 3B). These data lend strong support to our mapping results that identify the MxB targeting site to the tri-hexamer interface on capsid.

**Figure 3:**
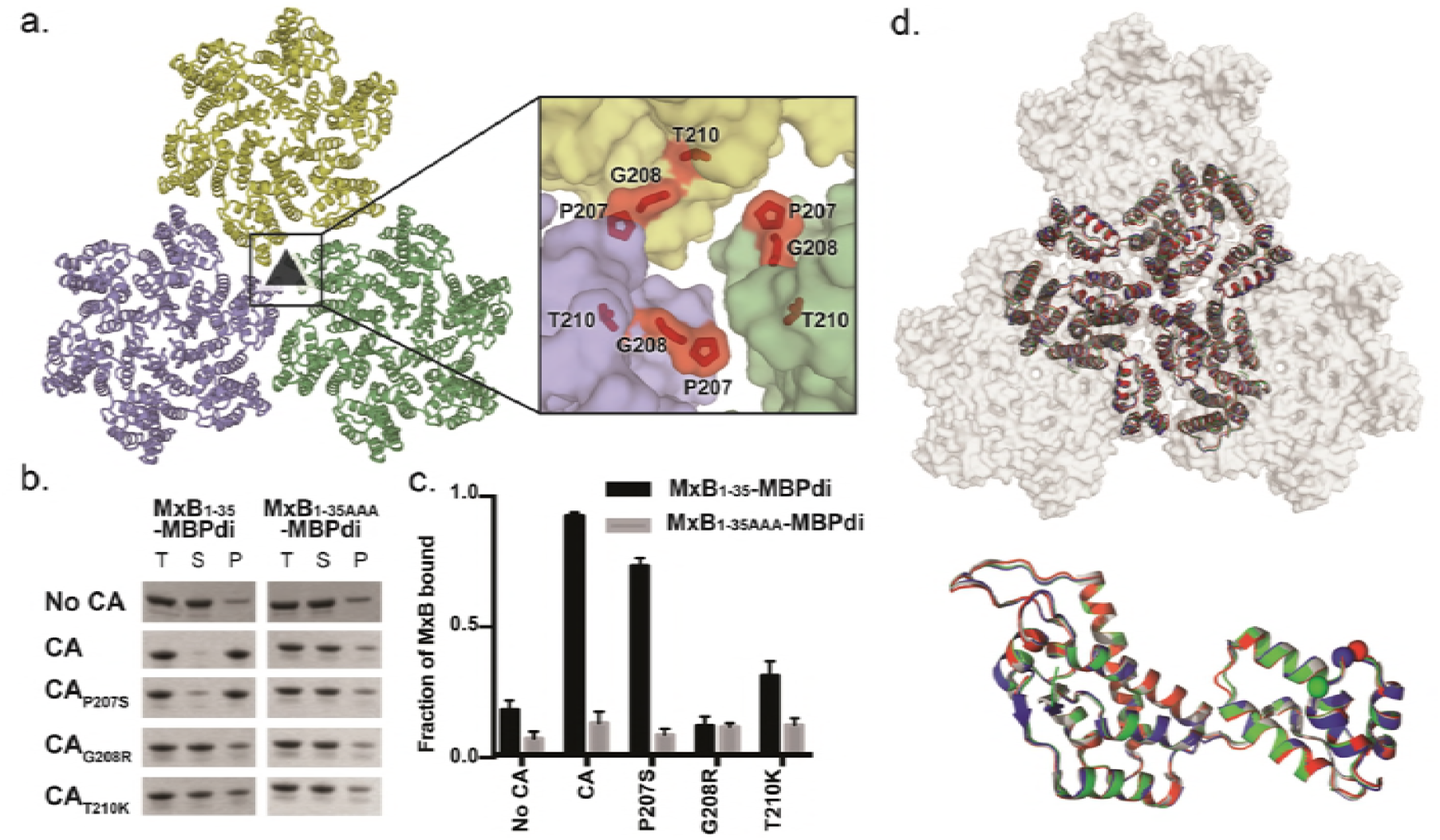
CA mutations that escape MxB restriction severely reduce MxB_1-35_-MBPdi binding. a. P207, G208, and T210 (red sticks) are located at the interface between three CA hexamers (light blue, light green, light yellow surface).
b. MxB_1-35_-MBPdi binding to CA tubes and CA tubes containing the mutations P207S, G208R in co-pelleting assays. The total, soluble, and pellet fractions were analyzed by SDS-PAGE. For simplicity, only the MxB_1-35_-MBPdi band is shown. Full gels and positive and negative controls performed with MBP-CCCyp and MBP, respectively, are shown in Supplemental Appendix 1.
c. Quantification of data in (b), performed as in figure 1. Error bars represent the standard error of the mean from three independent experiments. Quantification of the positive and negative controls can be found in Supplemental Appendix 1.
d. Alignment of P207S (red), G208R (blue), and T210K (green) crystal structures to the structure of WT CA (white, PDB:4XFX) at the tri-hexamer interface (top) and within a monomer (bottom). All of the mutations preserve the wild-type lattice structure, with only minor differences.

**Figure 4:**
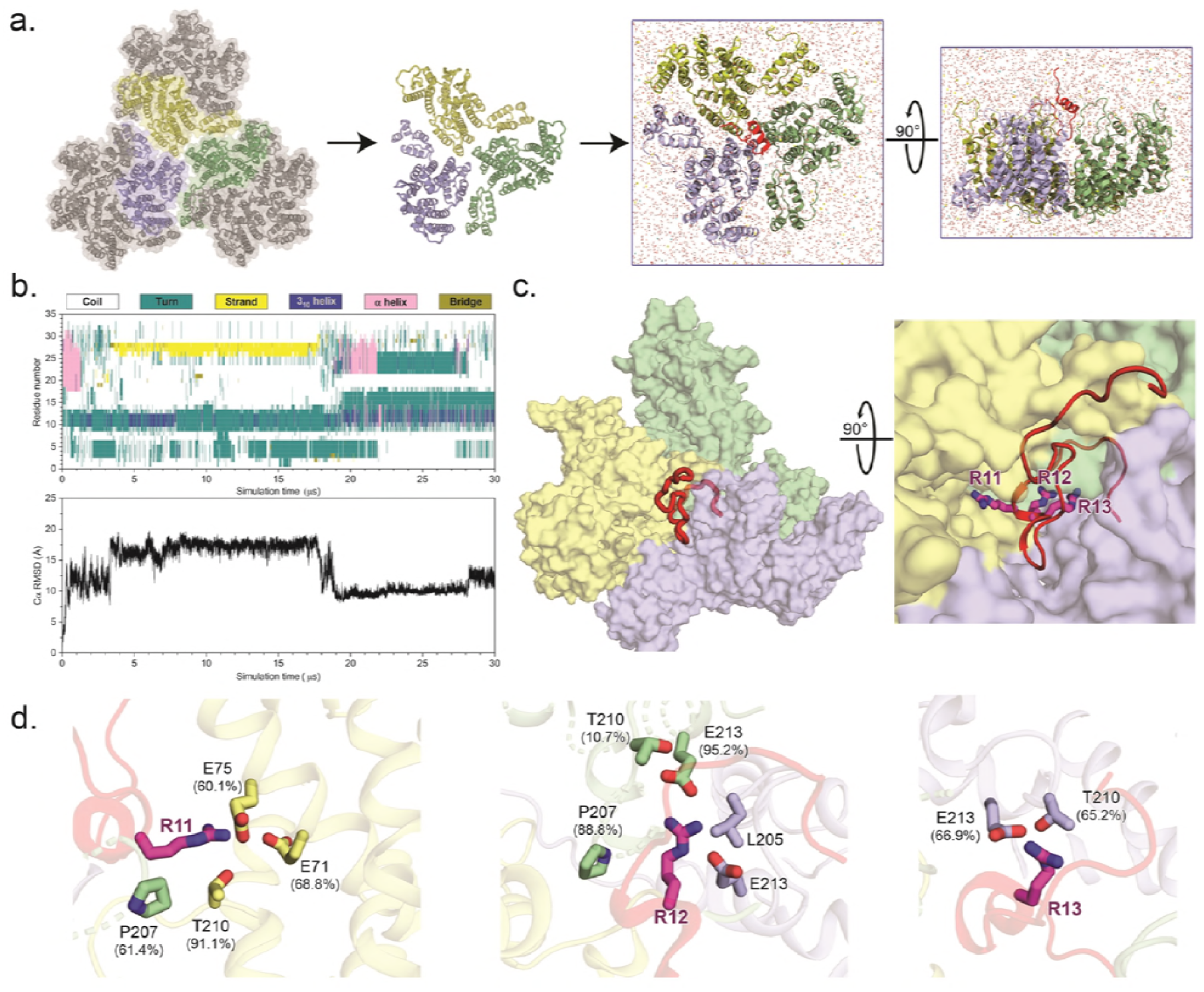
All-atom molecular simulations identify MxB interactions at the tri-hexamer interface. a. The all-atom model of MxB_1-35_ at tri-hexamer interface. Three dimers from different hexamers are in blue, yellow and green (left). The MxB_1-35_ peptide in red is placed above the middle of the tri-hexamer region before simulations (right).
b. The dynamics of MxB_1-35_ in tri-hexamer interface. The evolution of MxB secondary structure (top) during the simulation, computed by the STRIDE program in VMD and the root-mean-square deviation (RMSD) plot of the MxB Cα atoms (bottom).
c. MxB_1-35_ (red) binds in the tri-hexamer interface. The MxB ^11^RRR^13^ motif is shown in magenta sticks.
d. Molecular contacts between the MxB ^11^RRR^13^ motif and experimentally tested CA residues are shown for R11 (left), R12 (middle), and R13 (right), with corresponding contact occupancies.

### Hexamer interface mutations that affect MxB interaction do not change the conformation of CA lattice

We validated that the reduction in MxB binding observed for the hexamer interface mutations was due to a direct disruption in the MxB-capsid interface and not due to indirect changes in the CA lattice by examining the structure. Large conformational changes in the overall CA lattice could potentially lead to the loss of MxB binding at other sites. We first validated the overall morphology of the CA mutants tested by examining crosslinked CA tubes with negative stain electron microscopy. Morphologically, all CA mutant tubes appear similar to crosslinked tubes without mutation (SI Appendix 2). We further determined the crystal structures of those CA mutants that affect MxB restriction – P207S, G208R, and T210K – at resolutions ranging from 3.0Å to 3.3Å (Table 1). These mutants crystalized in the same crystal form as that of the WT CA, where the crystal packing recapitulates the CA lattice on the viral capsid [25]. The crystal structures of the mutants closely align to the structure of WT CA, both at the individual CA monomer level (RMSD 0.2-0.3 Å) and lattice level (RMSD 0.3-1.2 Å) for the six CA molecules surrounding 3-fold axis between three hexamers) (Fig 3D and E). These results are consistent with the idea that mutations at the tri-hexamer interface directly disrupt MxB-binding.

**Table 1:** Crystal structures of CA mutants P207S, G208R, T210K.

|  | **P207S** | **G208R** | **T210K** |
| --- | --- | --- | --- |
| PDB ID: | 6MQA | 6MQO | 6MQP |
| **Data Collection** |  |  |  |
| Space group | P6 | P6 | P6 |
| Cell dimensions |  |  |  |
| a, b, c | 90.34, 90.34, 57.62 | 92.01, 92.01, 57.19 | 92.04, 92.04, 57.35 |
| $\alpha, \beta, \gamma$ | 90, 90, 120 | 90, 90, 120 | 90, 90, 120 |
| Molecules/ASU | 1 | 1 | 1 |
| Resolution (Å) | 3.20 (3.39-3.20) | 3.00 (3.18-3.00) | 3.30 (3.49-3.30) |
| $R_{merge}$ | 12.8 | 10.4 | 13.9 |
| $I/\sigma$ | 6.6 (1.4) | 10.7 (2.7) | 7.6 (1.7) |
| Completeness (%) | 98.9 (99.1) | 98.3 (97.3) | 99.3 (99.4) |
| Redundancy | 2.6 | 3.1 | 3.5 |
| Unique reflections | 8591 (1377) | 8809 (1407) | 8116 (1320) |
| **Refinement** |  |  |  |
| # of non-hydrogen atoms | 1666 | 1693 | 1644 |
| R<sub>work</sub>/R<sub>free</sub> (%) | 18.6/21.9 | 20.6/27.1 | 20.7/24.0 |
| Average B factor | 73.6 | 76.0 | 87.6 |
| Root mean-squared deviation (rmsd) |  |  |  |
| Bond lengths | 0.005 | 0.017 | 0.01 |
| Bond angles | 1.0 | 1.8 | 1.1 |
| Ramachandran analysis |  |  |  |
| Preferred regions (%) | 97.14 | 97.64 | 98.11 |
| Allowed regions (%) | 2.86 | 2.36 | 1.42 |
| Outliers (%) | 0 | 0 | 0.47 |

Data collection and refinement statistics for CA<sub>P207S</sub>, CA<sub>G208R</sub>, CA<sub>T210K</sub>. Values shown in parentheses are for highest-resolution shell.

During purification, we observe that CA_G208R_ and CA_T210K_ formed hyperstable CA assemblies. These proteins have substantially lower solubility and visibly polymerize upon concentration and freeze/thaw. We believe this is due to the introduction of positive charge at a negatively-charged region of the CA lattice, resulting in charge neutralization that increases the stability of the CA lattice. In wild-type CA, it is likely that ideal lattice stability is maintained through balanced repulsion of negative charges at this interface. However, in the G208R and T210K mutants, the total negative charge at this interface is reduced, resulting in less lattice destabilizing repulsion force. A hyperstable capsid deviating from a fine balance could have deleterious effects for the virus, which may explain why these mutations are rarely seen in patient samples [26].

### Molecular dynamics simulations identify the tri-hexamer interface as the site of MxB-CA interaction

To investigate how MxB targets CA at an atomistic level, we constructed a CA tri-hexamer interface model and placed the 35 N-terminal residues of MxB (MxB_1-35_) in close proximity (Fig 4A). The model is composed of three dimers from adjacent CA hexamers, a structural motif commonly referred as a trimer-of-dimers. Subsequently, the model was subjected to thirty microseconds of all-atom MD simulation without any experimental restraints (SI Video 1). The canonical MD simulations permit MxB’s N-terminus and CA trimer-of-dimers to sample several physical conformations.

Results from our MD analysis show that MxB_1-35_ binds persistently in the tri-hexamer region throughout the entirety of the simulation, with extensive molecular contacts between MxB N-terminal region and CA residues (Table 2). Importantly, CA residues in the tri-hexamer interface, E71, E75 and E213, interact with the ^11^RRR^13^ motif of MxB with high occupancies (Fig 4D and Table 2), which is in good agreement with our co-pelleting assay data (Fig 2C and 2D). In fact, there is a remarkable agreement between the MD results with the experimental binding data obtained independently. For example, contacts between E212 and the MxB N-terminus were observed less frequently, which is consistent with the experimental finding that the E212A mutation has a smaller disruption of MxB binding than other glutamate mutations do at the tri-hexamer interface (Fig 2D). Moreover, we observe CA residues T210, G208, and P207 interacting with the ^11^RRR^13^ motif of MxB, which accounts for the importance of these sites in MxB binding (Fig 3B) and sensitivity [9]. In addition, we found contacts between CA and residues outside of the ^11^RRR^13^ motif (Table 2, SI Appendix 3), for example the cation-π interaction between CA R82 and MxB W8 residue, which may contribute to the residual level of binding observed with MxB_1-35AAA_-MBPdi.

**Table 2:** Top 20 MxB-CA contacts from all-atom MD simulation.

| # | MxB residue | CA residue | Occupancy (%) |
| --- | --- | --- | --- |
| 1 | ARG12 | GLU213 | 95.2 |
| 2 | TRP8 | ARG82 | 94.1 |
| 3 | ARG11 | THR210 | 91.1 |
| 4 | TYR10 | GLU75 | 90.6 |
| 5 | ARG12 | PRO207 | 88.8 |
| 6 | ARG12 | LEU205 | 87.4 |
| 7 | ARG12 | GLY208 | 84.5 |
| 8 | TYR10 | ALA78 | 83.9 |
| 9 | ARG11 | GLY208 | 74.8 |
| 10 | ARG11 | GLU71 | 68.8 |
| 11 | LYS6 | THR210 | 68.5 |
| 12 | ARG13 | GLU213 | 66.9 |
| 13 | ARG13 | THR210 | 65.2 |
| 14 | LYS6 | GLU75 | 63.5 |
| 15 | ARG11 | PRO207 | 61.4 |
| 16 | ARG11 | GLU75 | 60.1 |
| 17 | PRO9 | GLY208 | 59.6 |
| 18 | ARG12 | ALA209 | 59.6 |
| 19 | PRO7 | THR210 | 56.8 |
| 20 | HIS5 | ALA78 | 53.0 |

MxB and CA residue that were experimentally mutated are shown in blue and red, respectively.

The MxB-binding pocket revealed by the simulation is primarily formed by key CA residues located in adjacent CA hexamers (Table 2 and Fig 4D and SI Appendix 3). The MxB ^11^RRR^13^ motif is anchored to the bottom of the pocket, primarily interacting with the negatively-charged CA E213 residues, while the rest of the MxB_1-35_ peptide exhibits significantly higher flexibility (Fig 4D and SI Video 1). This observation is supported by the experimental finding that the E213A mutation shows the largest effect on MxB binding (Fig 2D). Notably, the significant difference in effect between E212A and E213A on MxB binding is consistent experimentally and computationally. This is likely due to their relative positions at the interface: E213 is closer to the 3-fold symmetry axis (Fig 2b) and other E213 residues from adjacent hexamers, such that MxB can bind E213 from multiple CA monomers simultaneously (Fig 4D). Bioinformatics tools predict the MxB N-terminal region to be disordered by itself, and it is either disordered or not present in the available 3D structures of MxB [11, 27]. Our secondary structure analysis (Fig 4B) shows that while the bound MxB_1-35_ still contains large disordered areas, the region near ^11^RRR^13^ motif maintains a stable conformation during the simulation (SI Video 1), which further supports the importance of the ^11^RRR^13^ motif in MxB-capsid interaction.

In addition, the MD simulation provides insight into the structure of the MxB_1-35_ upon binding to capsid and its influence on the ionic environment. The simulation shows that binding of MxB_1-35_ in the tri-hexamer interface decreases the sodium ion occupancies in the CA lattice. Based on the trajectory analysis, when the MxB N-terminal region occupies one of the three grooves between hexamers at the tri-hexamer interface, the calculated occupancy of sodium ions in the groove proximal to the bound region are lower than corresponding regions in the other two unoccupied grooves (SI Appendix 4). On the other hand, the chloride ion occupancy does not show appreciable difference in three grooves. Together with the high-occupancy molecular contacts between the positively charged ^11^RRR^13^ and the CA glutamates in the tri-hexamer region, these data agree with the experimental observation that MxB binding is salt-dependent, the interaction may lead to displacement of sodium ions at both the binding interface and nearby regions.

## Discussion

MxB is a potent restriction factor of HIV-1 infection. It restricts the nuclear important step of the viral life cycle. The inhibition requires an interaction with the HIV-1 capsid. Previous studies suggested that MxB recognized features present only in assembled capsid: interfaces between CA hexamers forming a lattice. Despite extensive work to identify CA mutations which affect MxB activity [5-7, 9, 11, 13, 18, 28], the interaction site and mechanism of interaction remain unclear. While cellular assays are efficient ways to identify mutations, they often fall short of disentangling the pleomorphic effects of CA mutations, where a functional defect in MxB sensitivity may not be directly related to MxB binding to capsid. Instead, it could be due to altered capsid stability or affecting other cellular factors that are also important in the MxB restriction pathway. Our biochemical and biophysical experimental system provides a stringent test to identify and verify mutations that have a direct effect on the MxB-capsid interaction. Combined with all-atom molecular dynamic simulations, our results demonstrate the molecular basis of how capsid mutations evade MxB restriction and provide insight into how MxB recognizes the capsid (Fig 5A).

**Figure 5:**
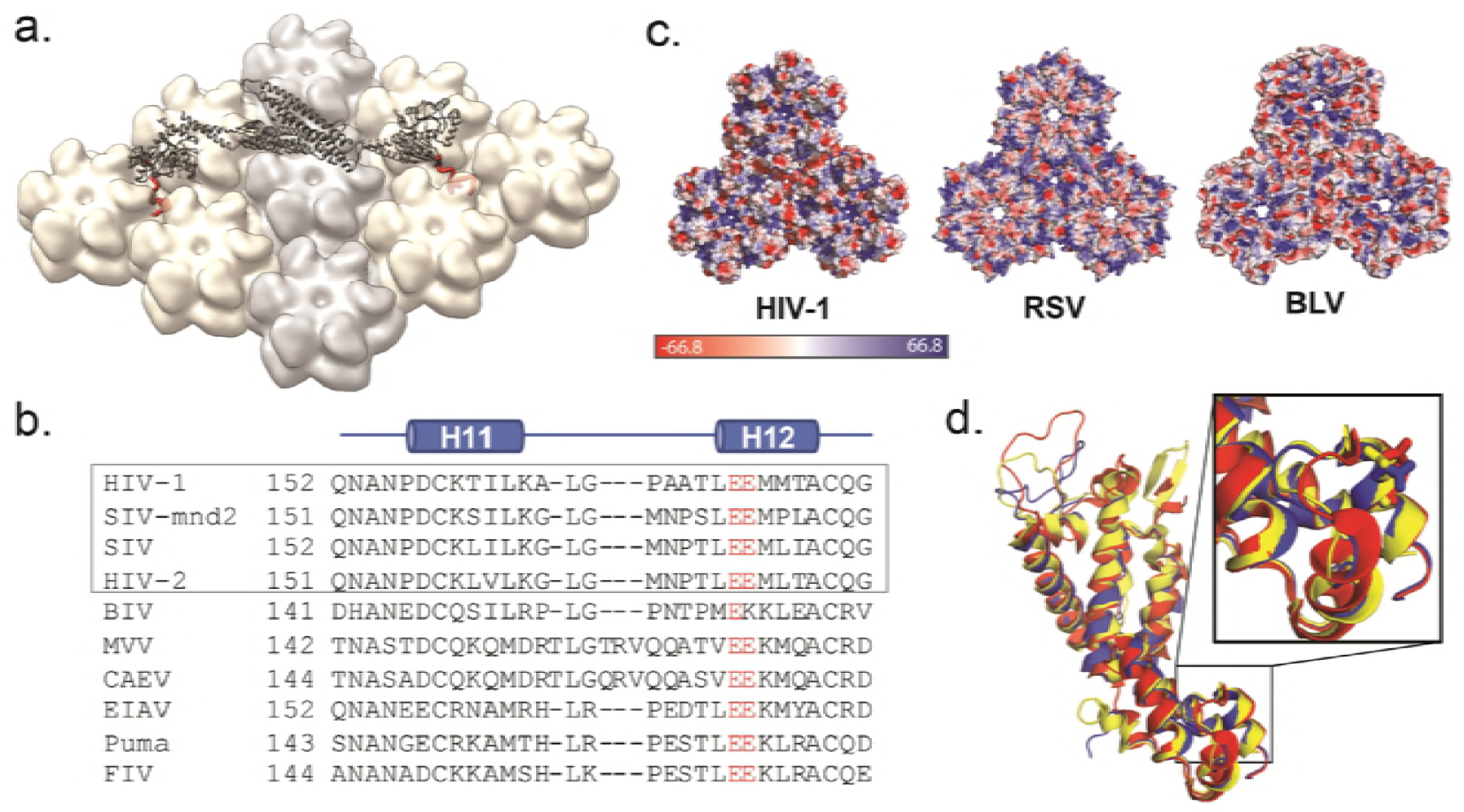
MxB recognizes CA features that are potentially conserved in lentiviruses but not in retroviruses. a. Model of MxB binding to the CA lattice, generated using existing structural data of the MxB dimer (gray, PDB ID: 4WHJ) and HIV CA (light gray/tan, PDB ID: 4XFX) and MD simulation data of the MxB N-terminal peptide binding to CA (red).
b. Alignment of CA structures available from different lentiviruses: HIV (PDB:4XFX, red, EIAV (PDB ID: 2EIA, blue), FIV, PDB ID: 5NA2, yellow); residue E213 shown in sticks located in the same position in each CA.
c. CA protein sequence alignment of lentiviruses. Gray box _denotes_ primate lentiviruses, which are restricted by MxB. Residue E213 and its homologous residues (red) are conserved in the lentiviral family.
d. Surface electrostatic potential distributions of diverse retroviral capsids show few shared characteristics at the tri-hexamer interface. The following structures are use in comparison: HIV-1 (PDB ID: 4XFX), RSV (PDB ID: 4TIR), BLV (PDB ID: 4PH0).

Our results paint a clear picture of MxB targeting HIV capsid at the interface between CA hexamers. To map the site of the MxB-capsid interaction, we performed systematic alanine scanning guided by an analysis of the electrostatic surface of the CA lattice, as well as previously reported MxB-escape mutations [9, 29, 30]. Mutations of the negatively charge residues surrounding the 3-fold axis on capsid, including E71/E75/E212/E213, had substantial deleterious effect on MxB interaction. These residues line a negatively charge well at the tri-hexamer interface, which is likely the binding site for the positively charged ^11^RRR^13^ motif in MxB that is critical for capsid interaction and HIV-1 restriction [14]. Consistently, we found that the tri-hexamer interface mutations that escape MxB restriction, G208R and T210K, also significantly disrupted the binding of MxB, likely due to a combined result of steric clash and charge repulsion with the ^11^RRR^13^ motif. In contrast, mutations in the di-hexamer interface and around the CypA-binding loop on the hexamer surface had little effect on MxB binding. Our structural studies showed these mutations had no apparent effect on the structure of CA lattice assemblies, supporting that the mutagenesis results revealed a direct binding of MxB at the tri-hexamer interface.

Though our mutant CA tubes are similar in morphology to that of wild-type CA, many of these mutations have deleterious effects in the virus, which may explain why MxB escape mutations are not commonly seen. Notably, it has been reported that the mutation E213A leads to decreased infectivity in cellular assays, and while Gag processing and viral cores appear normal, the core remains stable longer than wild-type upon entry into the target cell [31]. This is consistent with our observation during protein purification that E213A appeared to produce more stable CA assemblies and prone to precipitation. Viruses containing the CA mutations E71A and E180A had reduced infectivity relative to wild-type [32, 33]. The latter suggests that these two mutations are not tolerated well by the virus, and perhaps coping with MxB restriction is a lower-cost measure than evolving resistance.

Our all-atom molecular dynamics simulations identified the same important tri-hexamer interface independent of the experimental biochemistry. In the simulation, the bound MxB peptide interacts with many CA residues that were tested biochemically to be critical for MxB interaction (E71, E75, E212 and E213) and shown to be functionally important for MxB sensitivity (P207, G208, and T210). The extensive contacts between the MxB peptide and CA residues illustrate where and how MxB targets the CA lattice. Moreover, the glutamates in the tri-hexamer region, i.e. E213, create inter-hexamer repulsion and undermine the tri-hexamer interface structure and thus the CA lattice [31]. Therefore, these MxB-CA contacts imply that the MxB N-terminus may stabilize the CA lattice by screening charge repulsions between inter-hexamer glutamates, introducing additional contacts connecting neighboring hexamers and reinforcing the tri-hexamer region in CA lattice. The inter-hexamer interactions give the capsid structural plasticity, which affects its stability [25]. It is possible that CA stabilization is related to how MxB restricts HIV, as altering the capsid stability likely disrupts the timing of uncoating and capsid-modulated processes such as nuclear import. Further study is necessary to test this hypothesis.

The interface recognized by MxB appears to be a conserved feature of lentiviral capsid structure. Structure-based sequence alignments show that the residues that make up the acidic grooves of the HIV capsid are conserved among lentiviruses, but not retroviruses more generally. Both Bovine Leukemia Virus (BLV) and Rous Sarcoma Virus (RSV), two divergent retroviruses not restricted by MxB, have surface charge and geometry drastically different from those of HIV [34, 35]. In the lentiviral family, available structures of Equine Infectious Anemia Virus (EIAV) and Feline Immunodeficiency Virus (FIV) capsid overlay well with the wild-type HIV capsid structure (Fig 5B) [36, 37]. Though non-primate lentiviruses appear to have the necessary capsid surface for MxB biding, they are not restricted by MxB, perhaps because they use different nuclear import pathways [38]. These observations suggest that the presence of a negatively-charged lattice interface may be necessary for MxB restriction, but that restriction may be dependent on other factors like nuclear import pathways.

The work presented herein advances the mechanistic understanding of MxB restriction in molecular detail. We defined the HIV-1 capsid site recognized by the MxB ^11^RRR^13^ motif to be the interface between three CA hexamers. This represents a residue-level mapping of a HIV-1 capsid lattice-sensing restriction factor. Further work is needed to structurally characterize the interaction of MxB N-terminus and its interaction with CA. In addition, a better understanding of the binding affinity and behavior of the MxB oligomer will shed more light on its restriction. Due to the conservation of this interface among retroviruses, the tri-hexamer interface may represent a general binding site for host factors and a fruitful target for novel therapeutics.

## Materials and methods

### MxB_1-35_-MBP expression and purification

Plasmids encoding MxB_1-35_-MBPdi and MxB_1-35AAA_-MBPdi were transformed in BL21(DE3) cells and grown in TB at 37C to an OD600 of 0.4-0.6. Flasks were cooled, and 0.5mM IPTG was added to express the MxB-MBP protein at 18C overnight. Cells were centrifuged and pellets were resuspended in 40mL lysis buffer (50mM Tris pH8, 300mM NaCl, 0.2mM TCEP) with one cOmplete EDTA-free protease inhibitor cocktail (Sigma) and lysed using a microfluidizer. Lysate was cleared (13,500RPM in an SS-34 rotor at 4C for 35 min) and loaded onto a 10mL gravity column of Ni-NTA agarose beads equilibrated in lysis buffer. The column was washed with 3 volumes of lysis buffer, and protein was eluted in 30mL of Nickel B (50mM Tris pH8, 300mM NaCl, 400mM Imidazole, 0.2mM TCEP). The elute was concentrated to a volume of 5-10mL and diluted 10-fold with buffer S_A_ (50mM HEPES pH7, 0.2mM TCEP). This was loaded onto a 5mL Separhose Q HP columns on an FPLC and eluted using a gradient of buffer S_B_ (S_A_ + 1M NaCl). Peak fractions were concentrated to a volume of <2mL and loaded onto a size exclusion column (S200PG) equilibrated in 50mM Tris pH 8 and 300mM NaCl. Peak fractions were collected, concentrated, and flash frozen in liquid nitrogen. Purity of the sample was checked by SDS-PAGE at each step. At each step, phenylmethane sulfonyl fluoride (PMSF) was added to a final concentration of 1mM to reduce proteolysis.

### CA expression and purification

To improve solubility, we expressed all variants of our 14C/45C/WM proteins as CA-Mpro-MBP-6xHis fusions, with a SARS Mpro protease cleavage site and 6xHis tag to aid in purification. BL21(DE3) cells containing the plasmid encoding variants of CA_14C/45C_-mpro-MBP6xHis were grown in Terrific Broth at 37°C to an OD_600_ of 0.4-0.6, then protein expression was induced with 0.5mM IPTG at 25°C and allowed to progress overnight. Cells were pelleted and pellets were resuspended in 40mL lysis buffer (50mM Tris pH8, 300mM NaCl, 0.2mM TCEP) and lysed using a microfluidizer. Lysate was cleared (13,500RPM in an SS-34 rotor at 4°C for 35 min) and loaded onto a 10mL gravity column of Ni-NTA agarose beads equilibrated in lysis buffer. The column was washed with 3 volumes of lysis buffer, and protein was eluted in 30mL of Nickel B buffer (lysis buffer + 400mM imidazole). The elute was concentrated to a volume of 5-10mL and diluted 10-fold with buffer Q_A_ (50mM Tris pH8, 0.2mM TCEP). This was loaded onto 2 stacked 5mL Separhose Q HP columns and eluted using a gradient of buffer Q_B_ (Q_A_ + 1M NaCl). All proteins eluted between 5 and 15% buffer Q_B_. Mpro protease [39] was added directly to the peak fractions and allowed to incubate at 4 degrees. After 2 days, this protein was diluted with buffer S_A_ (25mM HEPES pH 7, 0.2mM TCEP) and loaded onto Sepharose SP columns and eluted using a gradient of buffer S_B_ (S_A_ + 1M NaCl). Proteins eluted between 5-10% buffer S_B_. Peak fractions were concentrated to a protein concentration of 10-15mg/ml and dialyzed for 4 hours at 4°C into storage buffer (50mM Tris, 75mM NaCl, 30mM BME). Purity of sample was checked by SDS-PAGE at each purification step.

### Co-pelleting assay

CA tubes were assembled by dialysis for 2 nights at high salt (50mM Tris pH 8, 1M NaCl) and subsequently one night at no salt (50mM Tris pH8). Concentrations were normalized through quantification of the CA bands on SAD-PAGE gels. Co-pelleting assays were performed with 3uM of MxB_1-35_-MBPdi or control proteins, and 100uM CA. Stock solutions of capsid tubes (200uM) and MxB_1-35_-MBPdi, MBP, or CC-Cyp proteins (6uM) were prepared. CA tubes were diluted in CA buffer (50mM Tris pH 8). Test proteins were diluted with twice the final concentration of salt in buffer containing 50mM Tris pH 8 and 150/300/600mM NaCl. Samples were prepared in 1.5mL Eppendorf tubes by combining 24ul of MxB_1-35_-MBPdi or control protein and 24ul of CA (or 50mM Tris pH8 for CA-controls) and incubated for 30 minutes at room temperature on a Nutator mixer. After 30 minutes, a 14ul “total” sample was removed and the remaining 28ul were centrifuged for 15 minutes at 4°C, 20,000RPM in an SS-34 rotor. The soluble fraction was subsequently removed and the pellet resuspended in 28ul of 50mM Tris pH8. Gel samples of the soluble fraction were made by taking 14ul of the soluble fraction and adding loading dye. 4ul of each gel sample was analyzed by SDS-PAGE (Invitrogen) and stained using SimplyBlue stain (Thermo Fisher). ImageStudioLite was used to quantify proteins of interest in the total and soluble fractions. The fraction of protein bound to CA was calculated by subtracting the ratio of soluble to total protein of interest from 1. The data were graphed with standard deviation using Prism 7.

### Molecular dynamics (MD) simulations

The 3D structure of first 35 residues of MxB (MxB_1-35_) was built de novo in Rosetta3 [40] and then equilibrated in a water box for 500 ns. The structure of native HIV-1 CA monomer (PDB:4XFX) was used to build the CA tri-hexamer model (Fig 4A, B). The equilibrated MxB_1-35_ was initially placed above the tri-hexamer region. The combined structure was then solvated with TIP3P water [41] and neutralized by NaCl at 150 mM concentration. The resulting model had a total atom number of about 150,000 (Fig 4B), included protein, ions and water, in a simulation dimension of 125 x 121 x 138 Å^3^.

After model building, the systems were initially subjected to a minimization and thermalization step. During thermalization the system was heated from 50K to 310K in 20K increments. Subsequently, the whole systems were equilibrated for over 20 ns, while the carbon alpha of the peripheral helices of CA monomer and one end of MxB_1-35_ were restrained. The minimization, thermalization and equilibration steps were completed in NAMD2.12 [42].

This equilibrated model was then run on a special purpose computer Anton2 [43] in the Pittsburgh supercomputing center for a total simulation time of 30 μs. To mimic the CA hexamer lattice in the simulation, the Cα atoms of three sets of CA α-helices on the peripheries of the tri-hexamer model were applied with a harmonic restraint of 1 Kcal/mol Å^2^ in x, y and z directions. Also, the Cα atoms of MxB residue 35 was applied a harmonic restraint of 1 Kcal/mol Å^2^ at 40Å away from the COM (center of mass) of CA in z direction. CHARMM 36 [44] force field was employed for all simulations. During the simulation, the temperature (310 K) and pressure (1 atm) was maintained by employing the Multigrator integrator [45] and the simulation time step was set to 2.5 fs/step, with short-range forces evaluated at every time step, and long-range electrostatics evaluated at every second time step. Short-range non-bonded interactions were cut off at 17 Å; long range electrostatics were calculated using the k-Gaussian Split Ewald method [46].

### MD trajectory analysis

The trajectory analyses were performed in VMD [47]. The MxB-CA contacts and their occupany scores were calculated by a homemade Tcl script. The secondary structure of MxB and ion occupancies were calculated with VMD plugins TimeLine and VolMap Tool, respectively.

### Crystallization and crystal structure analysis

CA crystals were obtained using the microbatch under oil method [48]. Purified CA P207S, G208R, or T210K was diluted to 1mg/ml (P207S) or 0.5 mg/ml (G208R or T210K) in 50mM Tris pH 8 buffer. 1ul of protein solution was mixed with 1ul precipitant solution (15% PEG3350, 0.1M NaI, 0.1M buffer) under oil (2:1 parafin: silicon). P207S crystals were grown with BisTrisPropane pH 6.5 while G208R and T210K were grown with Sodium Cacodylate pH 6.6 and 5.8 respectively. Crystals were harvested after several weeks of growth, cryo-protected in paratone oil and flash frozen in liquid nitrogen. Data were collected at the Advanced Photon Source, NE-CAT beamline 24-ID-C. Data processing was performed in XDS [49, 50]. The structure of WT CA (PDB: 4XFX) was used as a model for molecular replacement with the CCP4 program Phaser [25, 51-53]. Iterative refinement and manual rebuilding were performed using Refmac5, Phenix, and Coot [54-56]. Molecular graphics were generated using PyMol [57] and the MxB-capsid model was generated using Chimera [58].

### Electron Microscopy

Assembled CA tubes were diluted to 10uM in 50mM Tris pH 8 and negative-stained with uranyl acetate onto holey carbon grids after glow-discharge. Grids were examined on an FEI Talos L120C microscope and images at 57,000X magnification were collected using TEM Imaging and Analysis software (Gatan).

## Acknowledgements

We thank the staff at the Advanced Photon Source beamline 24ID-C. Anton computing time was provided by the Pittsburgh Supercomputing Center (PSC) and the Anton machine was generously made available by D.E.Shaw Research.

## SI captions

**SI Appendix 1: Co-pelleting supplemental data**. Full SDS-PAGE gels of co-pelleting assays, including MBP and CC-Cyp controls their quantification.

**SI Appendix 2: Electron micrographs of negative stained CA tubes used in this study**

**SI Appendix 3: Additional molecular contacts**. Molecular contact figures between MxB residues, K6 (left), W8 (middle), and Y10 (right), and CA residues.

**SI Appendix 4: Ion occupancies**. The ion occupancies of sodium (top) and chloride (bottom) calculated from the MD trajectory.

**SI Movie-1: The 30 μs MD simulation of MxB_1-35_ interacting with the CA tri-hexamer**

